# Microglia Depletion leads to Increased Susceptibility to Ocular Hypertension-Dependent Glaucoma

**DOI:** 10.1101/2024.03.05.583529

**Authors:** Cory A. Diemler, Michael MacLean, Sarah E. Heuer, Amanda A. Hewes, Olivia J. Marola, Richard T. Libby, Gareth R. Howell

## Abstract

In recent years, microglia have been highlighted for playing integral roles in neurodegenerative diseases, like glaucoma. To better understand the role of microglia during chronic ocular hypertension, we depleted microglia from aged (9-12 months old) DBA/2J (D2) mice, which exhibit age-related increases in intraocular pressure, using a dietary CSF1R antagonist, PLX5622. Retinal ganglion cell (RGC) somas were counted, and optic nerve cross-sections stained and assessed for glaucomatous damage. Sustained administration of dietary PLX5622 significantly reduced the numbers of retinal microglia. Dietary PLX5622 did not lead to changes in intraocular pressure in D2 or normotensive DBA/2J-*Gpnmb^+^* (D2-*Gpnmb^+^*) control mice. While PLX5622-treated D2-*Gpnmb^+^* did not develop optic nerve damage, PLX5622-treated D2 mice showed a significant increase in moderate-to-severe optic nerve damage compared to D2 mice fed a control diet. In conclusion, global reduction of microglia exacerbated glaucomatous neurodegeneration in D2 mice suggesting microglia play an overall beneficial role in protecting from ocular hypertension associated RGC loss.

## Introduction

Glaucoma is a leading cause of irreversible blindness worldwide (Tham et al., 2014), characterized by the progressive death and dysfunction of retinal ganglion cells (RGCs), the projection neurons that form the optic nerve connecting the retina to the brain (Ramirez et al., 2017, Tribble et al., 2020, Quigley and Broman, 2006, Howell et al., 2011, Williams et al., 2019). Aging and elevated ocular hypertension (OHT; elevated intraocular pressure, IOP) are the two major risk factors associated with glaucoma (Ramirez et al., 2017, Tribble et al., 2020, Quigley and Broman, 2006, Howell et al., 2011, Williams et al., 2019). Current treatments for glaucoma focus on lowering IOP, but not all patients with glaucoma present with increased IOP, and treatments focusing on normalizing IOP do not always successfully prevent vision loss (Schuster et al., 2020). Even when individuals receive proper treatment, approximately 10% still experience vision loss (Quigley and Broman, 2006), underscoring the need to identify new therapeutic targets that protect and prevent the loss of RGC function.

Microglia in the retina and optic nerve influence the progression of glaucoma (Howell et al., 2012, Heuss et al., 2018, Tribble et al., 2020, Howell et al., 2011, Rashid et al., 2019, Williams et al., 2019, Harder et al., 2020). They are the resident immune cell of the central nervous system and have distinct sets of functions for maintaining homeostasis, including removal of cellular debris, monitoring for injury and pathogens and coordinating immune responses (Sun et al., 2021, Ramirez et al., 2017, Bosco et al., 2008, Heuss et al., 2018, Tribble et al., 2020, Rashid et al., 2019, Bachiller et al., 2018, Hickman et al., 2018, Song and Colonna, 2018, Bosco et al., 2012). Microglia survey and continually respond to their environment. In response to changes in the environment, such as an increase in IOP or injury, microglia adopt specific phenotypes which include morphological changes and adaptations to their transcriptional profiles. These changes are seen before RGC dysfunction and death (Williams et al., 2017b, Howell et al., 2011, Bosco et al., 2012). It is proposed that microglial dysfunction contributes to a sustained inflammatory environment leading to changes in retinal integrity and RGC loss (Rashid et al., 2019, Ramirez et al., 2017).

Previous work by others has shown that diminishing microglial activation via minocycline treatment (Bosco et al., 2008, Baptiste et al., 2005) or irradiation (Bosco et al., 2012) promoted RGC survival in DBA/2J (D2) mice, an aging OHT mouse model of glaucoma. However, both approaches likely affect a variety of other processes. A more specific approach to test the role of microglia is to deplete microglia using the colony stimulating factor 1 receptor (CSF1R) inhibitor PLX5622, which has been used to test the role of microglia in brain diseases such as Alzheimer’s disease (Spangenberg et al., 2019). The role of microglia in the eye has been studied in various diseases using PLX5622. In an optic nerve crush study, using PLX5622 revealed microglia are not critically involved in RGC degeneration after an acute neuronal injury (Hilla et al., 2017). In a diabetic retinopathy mouse model, PLX5622 treatment for approximately 2 months protected the retina from neurodegenerative and vascular damage (Church et al., 2022). Depletion of microglia using PLX5622 in a model of RGC degeneration, intravitreal injection of NMDA, led to RGC protection compared to controls (Takeda et al., 2018). In contrast, PLX5622 treatment in an acute inducible model of OHT resulted in increased RGC loss (Tan et al., 2022). However, widely used acute models of OHT have limited degeneration of RGCs and, given aging is the major risk factor for glaucoma, limited translational relevance. Therefore, to address this, we focused our efforts on D2 mice which develop glaucomatous neurodegeneration due to age-related elevation of IOP (Anderson et al., 2002). To study the role of microglia in D2 glaucoma, microglia were depleted using dietary PLX5622 beginning at 9.5 months of age, a time point after IOP elevation has begun and a timepoint when the vast majority of eyes have not developed glaucomatous neurodegeneration (Libby et al., 2005a). Glaucomatous neurodegeneration was assessed at 12 mos, when the majority of eyes show moderate to severe glaucoma (Libby et al., 2005a). Loss of microglia significantly increased the proportion of eyes with severe glaucomatous optic nerve damage and RGC soma loss, suggesting an overall beneficial role for microglia in the progression of glaucoma.

## Methods

### Ethics statement

All research was approved by the Institutional Animal Care and Use Committee (IACUC) at The Jackson Laboratory (JAX, approval number 12005). Animals were humanely euthanized using cervical dislocation. Authors performed euthanasia using methods approved by the American Veterinary Medical Association.

### Animals

All mice were housed in a 12-hour light/12-hour dark cycle under previously described conditions, and fed *ad libtum* (Howell et al., 2012, Howell et al., 2011). Our D2 and D2-*Gpnmb+* colonies are routinely crossed with D2 mice from JAX production facility to prevent genetic drift as previously described (Howell et al., 2011).

### Microglia depletion

Microglia were depleted by administering PLX5622, an inhibitor of the colony-stimulating factor 1 receptor, in the chow (Elmore et al., 2014). PLX5622 was acquired from Chemgood (C-1521) and formulated in Purina 5K52 mouse chow diet at a concentration of 1200mg/kg (ppm) by Research Diets Inc, followed by 10–20 kGy gamma irradiation. Chemical purity and proper diet concentration were validated through HPLC and mass spectrometry analysis through Chemgood and JAX Proteomics core. To evaluate efficiency of microglia depletion, 2-4 mos or 9.5 mos D2 mice were provided PLX5622 diet for 3 weeks. To evaluate the effects of microglia depletion on glaucoma-relevant outcomes, cohorts of 9.5 mos D2 or D2-*Gpnmb+* mice were provided PLX5622 diet for 10 weeks. Non-depleted control mice received standard 5K52 mouse chow diet. All cohorts included equal numbers of male and female mice.

### IOP measurements

To evaluate the effects of microglia depletion on IOP, a cohort of D2 mice were given dietary PLX5622 at 9.5 mos. At baseline and after 5 and 10 weeks of dietary PLX5622, IOPs were measured with a rebound tonometer (TonoLab; ICare, Raleigh, NC). Three readings were taken from each eye and averaged to obtain the reading for a timepoint.

### Retina and optic nerve isolation

At the appropriate age, all mice were euthanized. To obtain the retina, eyes were removed and fixed in 4% PFA overnight at 4°C. Thereafter, the cornea was cut along the limbus to remove the lens and vitreous and the retina and attached portion of the optic nerve head separated. Retinas were either processed immediately for myeloid cell analysis, snap frozen for RNA-seq, or stored in 1x Phosphate Buffered Saline (PBS) at 4°C for immunofluorescence. To obtain the retroorbital portion of the optic nerve for axon damage assessment, the head was separated from the body, the skull opened, and the top part of the brain removed. The head was then fixed in Smith Rudt buffer (0.8% PFA, 1.2% glutaraldehyde in 0.1M phosphate buffer) for 48 hours. Thereafter, the rest of the brain was removed to expose the retroorbital portion of the optic nerves (up to the optic chiasm). Left and right optic nerves were then carefully removed from the base of the skull and stored in 0.1M phosphate buffer at 4°C until required.

### Retina homogenization, myeloid cell preparation and flow cytometry analysis

Retinas were homogenized and pooled as previously described (Williams et al., 2017a). Eyes were enucleated and placed directly into ice-cold HBSS. Retinas were dissected and placed into 100uL of HBSS with dispase (5U/mL) DNAse 1 (2000U/mL). Retinas were incubated for 20 minutes at 37°C whilst being shaken at 350RPM in an Eppendorf Thermomixer R. Samples were gently triturated. For flow cytometric analysis, each sample was stained with DAPI, CD45 BV605 (clone 30-F11, BD Biosciences, 1:240), and CD11b PE (clone M1/70, Biolegend, 1:960), followed by processing on a FACSymphony A5 cytometer, and analysis using FlowJo (v10) software.

### RNA isolation, library preparation and RNA-sequencing

Retinas were lysed and homogenized in TRIzol Reagent (ThermoFisher) using a Pellet Pestle Motor (Kimbal), then RNA was isolated using the miRNeasy Micro kit (Qiagen), according to manufacturers’ protocols, including the optional DNase digest step. RNA concentration and quality were assessed using the Nanodrop 8000 spectrophotometer (Thermo Scientific) and the RNA 6000 Pico Assay (Agilent Technologies). Libraries were constructed using the KAPA mRNA HyperPrep Kit (Roche Sequencing and Life Science), according to the manufacturer’s protocol. Briefly, the protocol entails isolation of polyA containing mRNA using oligo-dT magnetic beads, RNA fragmentation, first and second strand cDNA synthesis, ligation of Illumina-specific adapters containing a unique barcode sequence for each library, and PCR amplification. The quality and concentration of the libraries were assessed using the D5000 ScreenTape (Agilent Technologies) and Qubit dsDNA HS Assay (ThermoFisher), respectively, according to the manufacturers’ instructions. Approximately, 70M 150bp paired-end reads were sequenced per sample on an Illumina NovaSeq 6000 using the S4 Reagent Kit v1.5 (Illumina, Cat#20028313) by the Genome Technologies Core at the Jackson Laboratory.

### Retina RNA-sequencing data analysis

Raw FASTQ files were processed using standard quality control practices (Wingett and Andrews, 2018). High quality read pairs were aligned to the mouse genome (mm10) using STAR 2.7.9a (Dobin et al., 2013). A custom genome was generated for D2 mice by incorporating REL-1505 variants into the reference genome (Keane et al., 2011, Dobin et al., 2013). Read pair counts were summed with the featureCounts function in subread 1.6.3 using the Ensembl Release 68 transcriptome reference (Liao et al., 2013, Cunningham et al., 2022). Differential expression analyses were completed using edgeR v3.40.2 within the R environment v4.2.3 using the glmQLFtest function (Robinson et al., 2010). Groups were defined as a combination of sex and treatment (∼0+Group). Contrasts were defined to test for the average treatment effect, the treatment effect within each sex, the average sex effect, and the interaction effect between sex and treatment. Genes were filtered using filterByExpr within edgeR prior to normalization. The clusterProfiler v4.6.2 R package was used to test for overrepresentation of KEGG gene sets within the list of differentially expressed genes (DEGs) with a false-discovery rate of less than 0.05 (Wu et al., 2021, Yu et al., 2012, Kanehisa and Goto, 2000).

### Analysis of glaucomatous damage

Intracranial portions of optic nerves were processed and analyzed as previously described (Libby et al., 2005b, Howell et al., 2011, Libby et al., 2005a, Anderson et al., 2005). Briefly, fixed optic nerves were processed and embedded in plastic. One-micrometer-thick cross sections of optic nerve from behind the orbit were cut and stained with paraphenylenediamine (PPD). Two observers blinded to the experimental protocol determined the degree of nerve damage using a validated grading method (Howell et al., 2007). No/early (NOE) damage was defined as less than 5% axons damaged. This amount of damage is seen in age-matched mice of various strains that do not develop glaucoma. It is called “No/early” because some of these eyes are undergoing early molecular stages of disease, but they are not distinguishable from eyes with no/early glaucoma by conventional morphologic analyses of axon number and nerve damage. Moderate (MOD) described nerves with many damaged axons throughout the nerve, with an average of 30% axon loss. Severe (SEV) nerves had substantial axon loss (>50% lost) and damage. The axon damage assessment scheme as previously been validated using axon counts (Libby et al., 2005b, Howell et al., 2011, Libby et al., 2005a, Anderson et al., 2005).

### Immunofluorescence and microscopy

Retinas were permeabilized with 1xPBS and 0.3% TritonX-100 (PBT), blocked for 24 hours at 4°C in PBT + 10% normal donkey serum (D9663-10ML, Sigma-Aldrich), washed once with PBT, and incubated in primary antibody solution containing goat anti-IBA1 (1:300, ab5076, Abcam) and rabbit anti-RBPMS (1:250, GTX118619, GeneTex). After primary incubation for 72 hours at 4°C, sections were washed 3 times with PBT and incubated in secondary antibodies (donkey anti-goat Alexa Fluor 568 and donkey anti-rabbit Alexa Fluor 488 (both diluted 1:1000 in PBT)) for 24 hours at 4°C. Retinas were washed with PBS and mounted. Images were captured using two methodologies. Retinas stained with anti-IBA1 were imaged on a Leica DM6 microscope at 20X magnification. For IBA1, 3 images per retina were captured at 1024 × 1024 pixel frames and counts were averaged. Analysis was performed using ImageJ2 (version 2.3.0/1.53q). Maximum projections were generated from stacks, individual channels were isolated, and default thresholds were applied for IBA1. Cell bodies were manually counted. Retinas stained with anti-RBPMS were imaged using a Leica Versa slide scanner at 10X magnification, capturing and merging individual tiles. RBPMS+ RGCs were counted across the whole retina. IMARIS software was used for analysis (v9.5.1) by using the RBPMS signal channel and quantifying RGC soma using Spots function (threshold soma size of 10 microns).

### Statistical analysis

Data were analyzed masked to treatment group. All statistical analyses were performed in GraphPad Prism9 software. Unless otherwise stated, data are presented as mean ± SEM. For IBA1+ microglia, Mann-Whitney test was performed for statistical significance because data failed Shapiro-Wilk test for normalcy. To determine differences in IOP within genotype/age/treatment, three-way ANOVA was performed for statistical significance. For RBPMS+ RGC counts, Kruskal-Wallis test was performed for statistical significance because data failed Shapiro-Wilk test for normalcy. Chi-squared tests were used to evaluate differences in optic nerve damage distributions among groups. Unless specifically stated in the text, *p* < 0.05 was considered significant.

## Results

### CSF1R inhibition causes significant retinal microglia depletion in young and aging DBA/2J mice

To evaluate microglia depletion efficiency in the retina, we first assessed retinal microglia numbers in 2-4 mos male and female D2 mice provided with PLX5622 diet (1200 ppm), a similar concentration to that used previously (Elmore et al., 2014). D2 mice given a standard chow diet were used as controls. After 3 weeks on PLX5622 diet, analysis of microglia number was determined by flow cytometry. A 75% reduction of CD45+ CD11b+ cells was observed in D2 retinas fed PLX5622 compared to the control diet (Figure1A). We next looked at the effects of three weeks of PLX5622 treatment on retinas from D2 mice starting at 9.5 mos (after OHT in the majority of D2 eyes) using RNA-seq. Compared to retinas from D2 mice on a control diet, retinas of D2 mice on PLX5622 showed many microglial-relevant genes were significantly downregulated (Figure 1B-C), and there was a significant enrichment of differentially expressed genes involved in microglia-relevant biological processes (Figure1D). Collectively, these data show that dietary PLX5622 administered for 3 weeks results in the majority of microglia being depleted from retinas of young and middle aged D2 mice.

**Figure 1.**
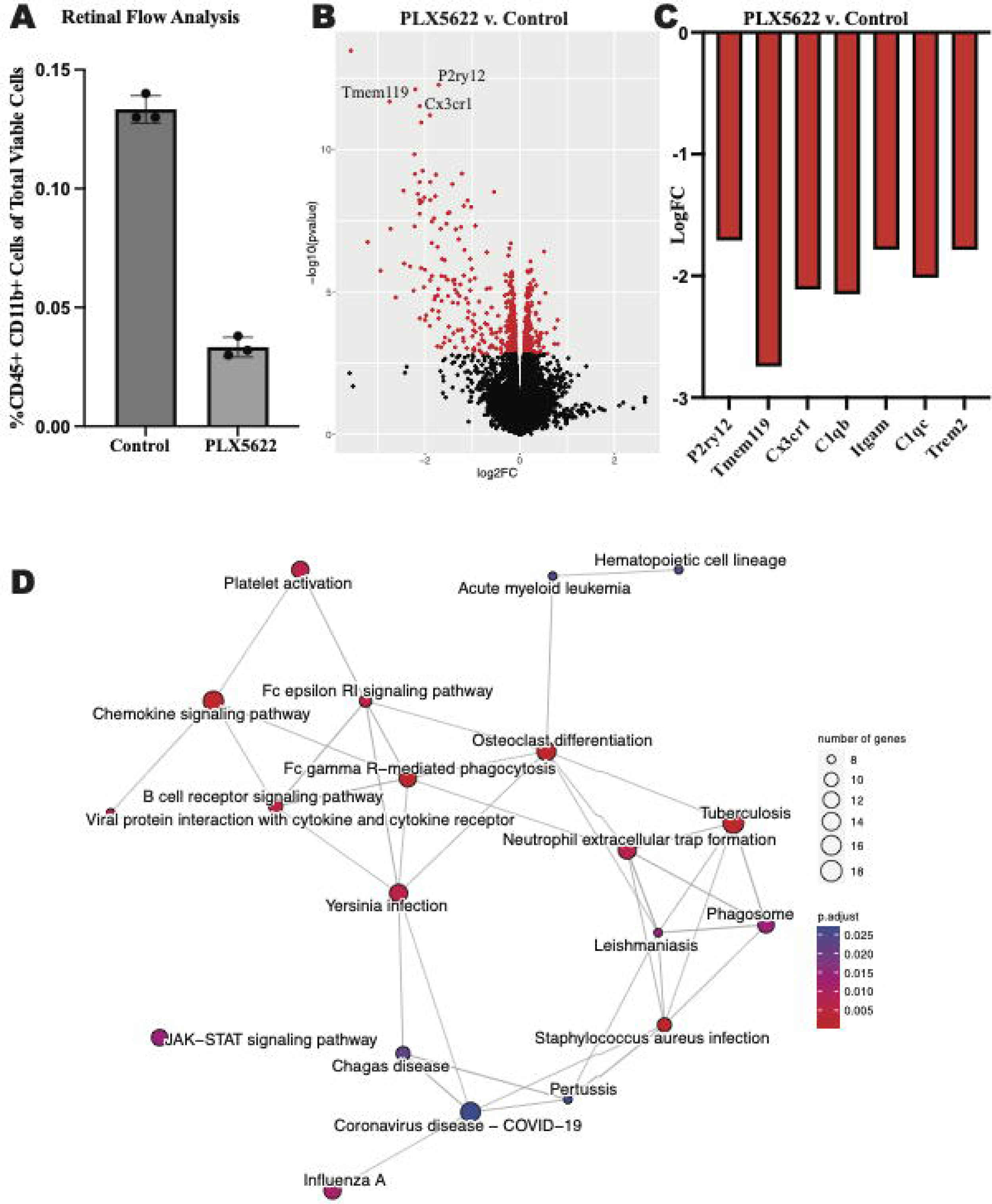
Dietary PLX5622 leads to retinal microglia depletion. **A.** Flow cytometry cell counts of CD45+ and CD11b+ cells between control diet (0.133±0.003) and 3 weeks of dietary PLX5622 (0.033±0.002) from 2-4 mos D2 retinas (data are presented as Mean±SEM, n=3; Welch’s t-test, p<0.0001; ) **B.** Volcano plot of RNA-seq results comparing retinas from 9.5 mos PLX5622 treated D2 compared to control diet D2. **C.** Expression levels of microglial-relevant genes were significantly down regulated. **D.** Enriched KEGG pathways in the DEGs identified between PLX5622 and control diet retinas illustrated as a network.

### CSF1R inhibition does not modify IOP levels or cause optic nerve damage in normotensive mice

To determine if PLX5622 treatment altered glaucoma-relevant phenotypes, D2 and normotensive D2*-Gpnmb^+^* controls (Howell et al., 2007) were provided either the PLX5622 diet or control diet from 9.5 to 12 mos. IOP was assessed at baseline and after 5 and 10 weeks of dietary PLX5622 (just prior to harvest). Dietary PLX5622 did not significantly alter the IOP of D2 or D2-*Gpnmb+* mice (Figure 2A). To determine whether PLX5622 treatment alone, independent of OHT, was sufficient to induce axon damage, optic nerves from D2-*Gpnmb*^+^ mice fed PLX6522 diet or control diet were assessed. Optic nerves from D2-*Gpnmb^+^* mice fed either PLX5622 diet or control diet showed no significant axon damage (Figure 2B-D), suggesting CSF1R inhibition, independent of OHT, is not sufficient to induce RGC loss.

**Figure 2.**
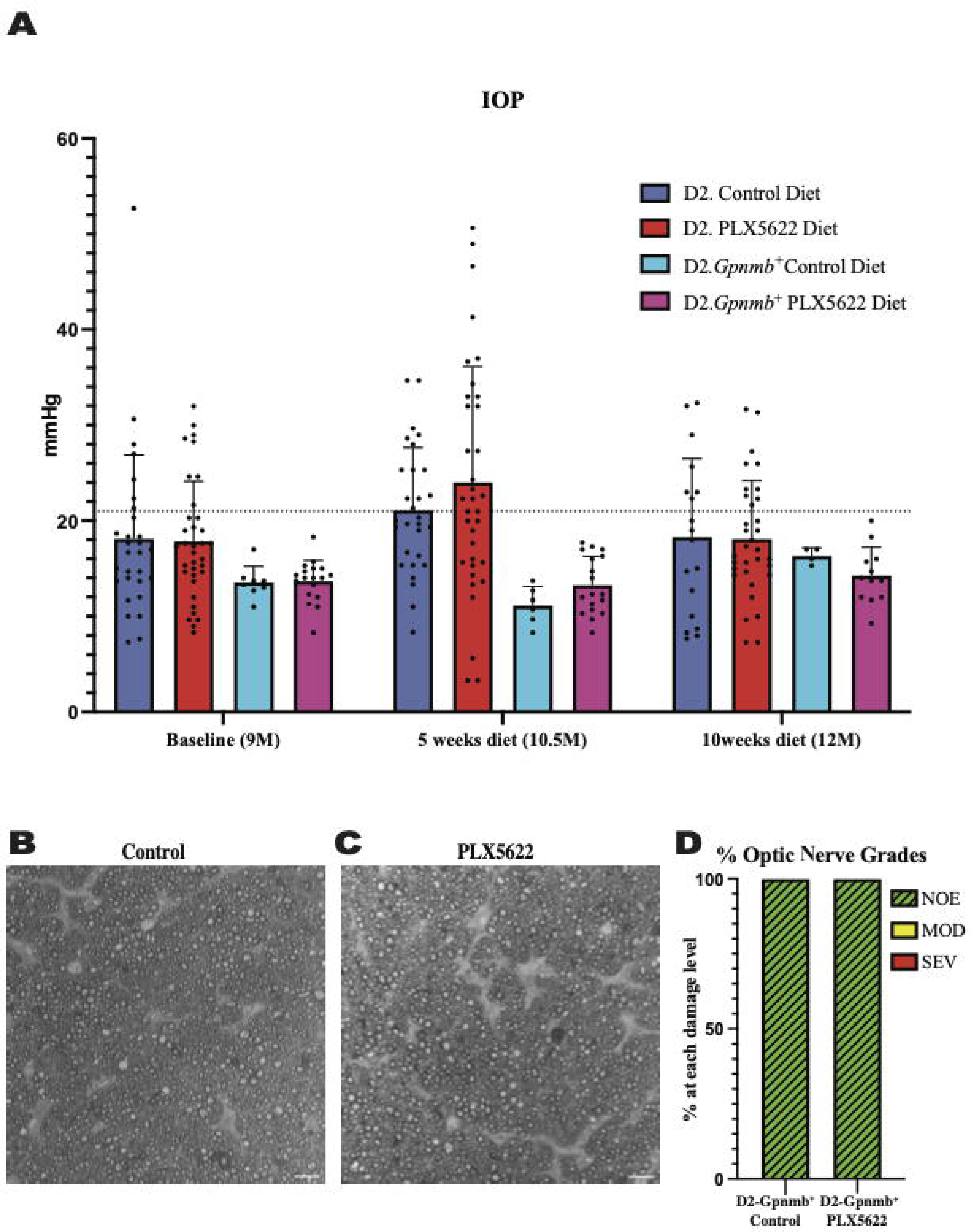
Microglia depletion does not alter IOP or optic nerve degeneration. **A.** IOP measurements of D2 mice on control diet, D2 mice on PLX5622 diet, D2-*Gpnmb+* mice on control diet, D2-*Gpnmb+* mice on PLX5622 diet were taken at 9mos (start of treatment), 10.5mos (5 weeks on diet) and 12mos (10 weeks on diet; data presented as Mean±SEM). **B-C.** Qualitative assessment of optic nerve health. RGC axons with and without CSF1R inhibition were stained with PPD, which differentially stains sick and dying axons. Examples of D2-*Gpnmb*^+^ optic nerves without dietary PLX5622 (**B**), and with dietary PLX5622 (**C**). **D.** The frequency of glaucomatous optic nerve damage in D2-*Gpnmb*^+^ mice on control diet (n=5) and PLX5622 diet (n=7) using a 3-point grading system (Methods). No difference in nerve damage was seen between treated and untreated eyes (Chi-square test, p<0.001).

### CSF1R inhibition increased the number of D2 eyes with glaucomatous neurodegeneration

To determine the effect of microglia depletion on glaucomatous neurodegeneration, D2 mice were provided dietary PLX5622 from 9.5 to 12 mos. Confirming our short-term administration studies (Figure 1), whole mount retinal immunofluorescence revealed a significant reduction in the number of IBA1, selective marker for microglia,+ cells in the inner plexiform layer (IPL) of PLX5622-treated mice compared to control chow mice (Figure 3). To determine changes in glaucoma severity, optic nerve cross-sections were evaluated for RGC axonal damage using PPD. In D2 mice fed the control diet, 74% of optic nerves showed moderate-to-severe glaucomatous neurodegeneration, similar to previous reports (Howell et al., 2012). In contrast, D2 mice fed dietary PLX5622, had a significant increase in glaucomatous damage, with 97% of optic nerves graded as moderate-to-severe (Figure 4).

**Figure 3.**
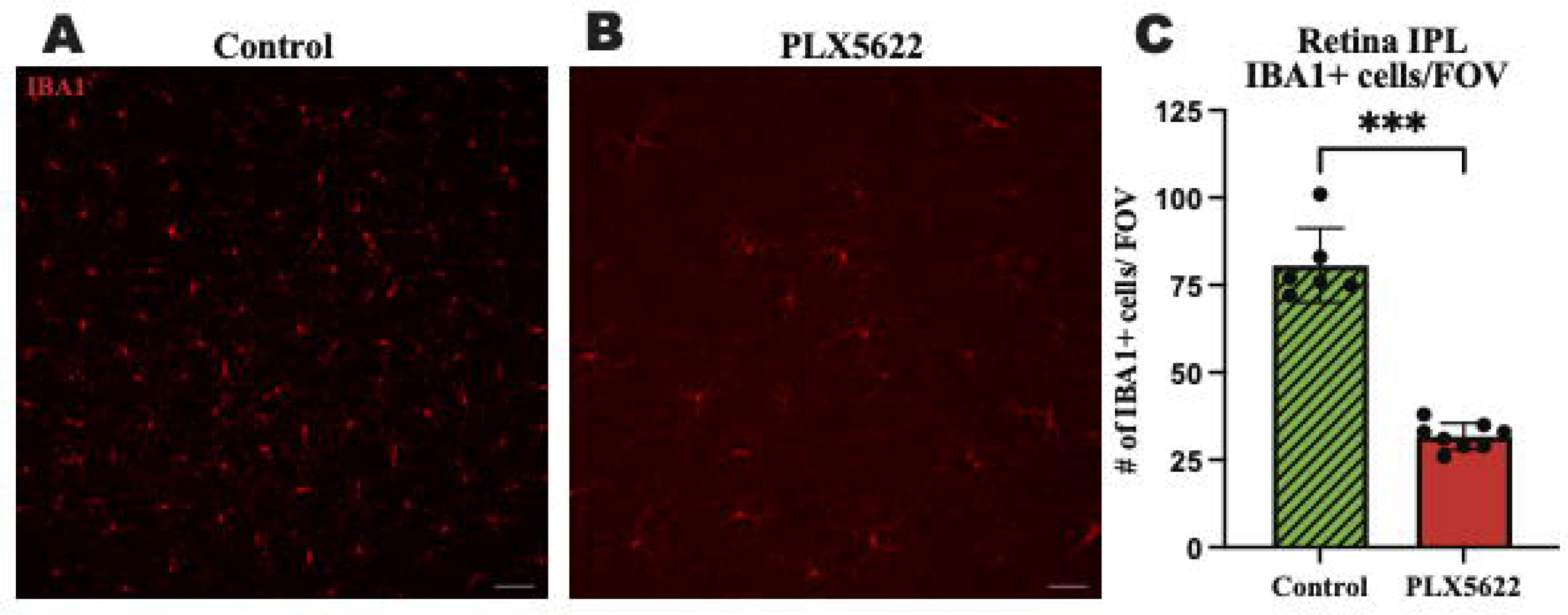
Prolonged CSF1R inhibition reduces microglia in aged D2 retinal tissue. **A-B**. Representative images of IBA1+ cells in the IPL of retinal flatmounts of 12mos D2 retinas with 10 weeks of **A.** control diet and **B.** PLX5622 treatment. **C.** Quantification of IBA1+ cells in the control diet (Mean±SEM, 80.7±10.6, n=6) and PLX5622 diet (31.8±3.8, n=8). Scale bar in **A** and **B**, 50µm; ****, p<0.0001 (Welch’s t-test).

**Figure 4.**
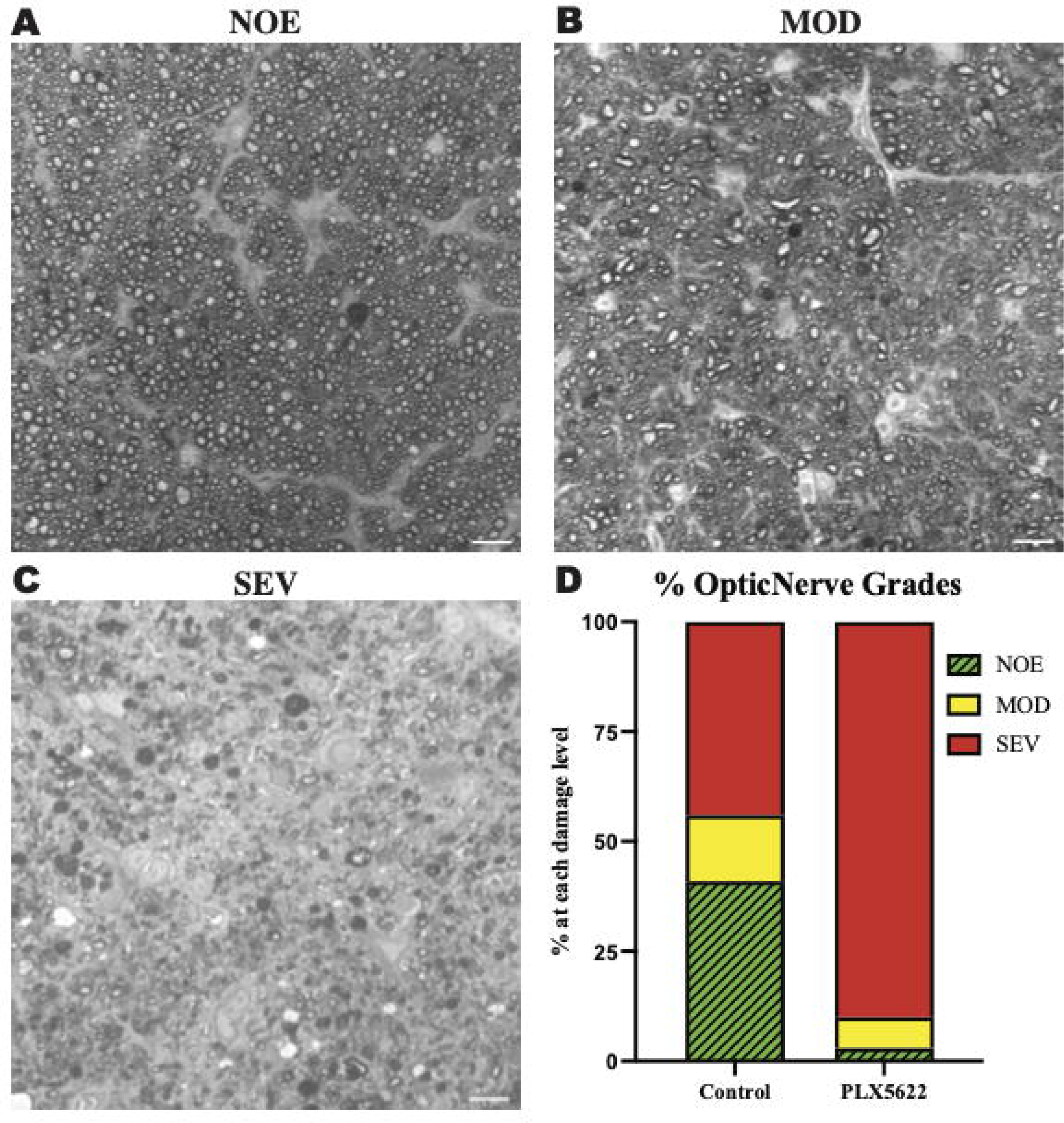
Microglial depletioin increased prevalence of severe optic nerve damage in D2 mice. **A-C.** Qualitative assessment of optic nerve health. RGC axons were stained with PPD, and examples of optic nerves with No/Early damage (NOE) (**A**), Moderate (MOD) (**B**), and severe (SEV) are shown (**C**). **D.** The frequency of glaucomatous optic nerve damage in D2 mice on control diet (n=13) and PLX5622 diet (n=17) using a 3-point grading system (Methods). There was an increased damage level observed in the PLX5622 treated eyes (p<0.0001, Chi-square test was performed). Scale bar in **A** and **B**, 10µm.

Previous studies using BAX-deficient D2 mice showed that the processes determining optic nerve degeneration could be uncoupled from RGC soma loss (Libby et al., 2005b). Therefore, it is possible that RGC somas are not lost in PLX5622-treated D2 mice with severe optic nerve damage. To test this, RBPMS+ RGCs from retinas from D2 mice fed the PLX5622 diet or control diet were counted (RBPMS is a marker of RGCs). Retinas from 12 mos D2 mice fed a control diet and classified as having no optic nerve damage, were used as the control. Retinas from D2 mice fed the control diet classified as having severe optic nerve damage showed a significant reduction in the number of RBPMS+ RGC somas compared to retinas with corresponding no/early damaged optic nerves. Retinas from D2 mice fed the PLX5622 diet classified as having severe optic nerve damage also showed a significant reduction in RBPMS+ RGC somas compared to no/early damaged optic nerve retinas (Figure 5). These data show that microglia depletion did not alter RGC soma survival and confirms that RGC axonal degeneration and RGC soma loss remain coupled.

**Figure 5.**
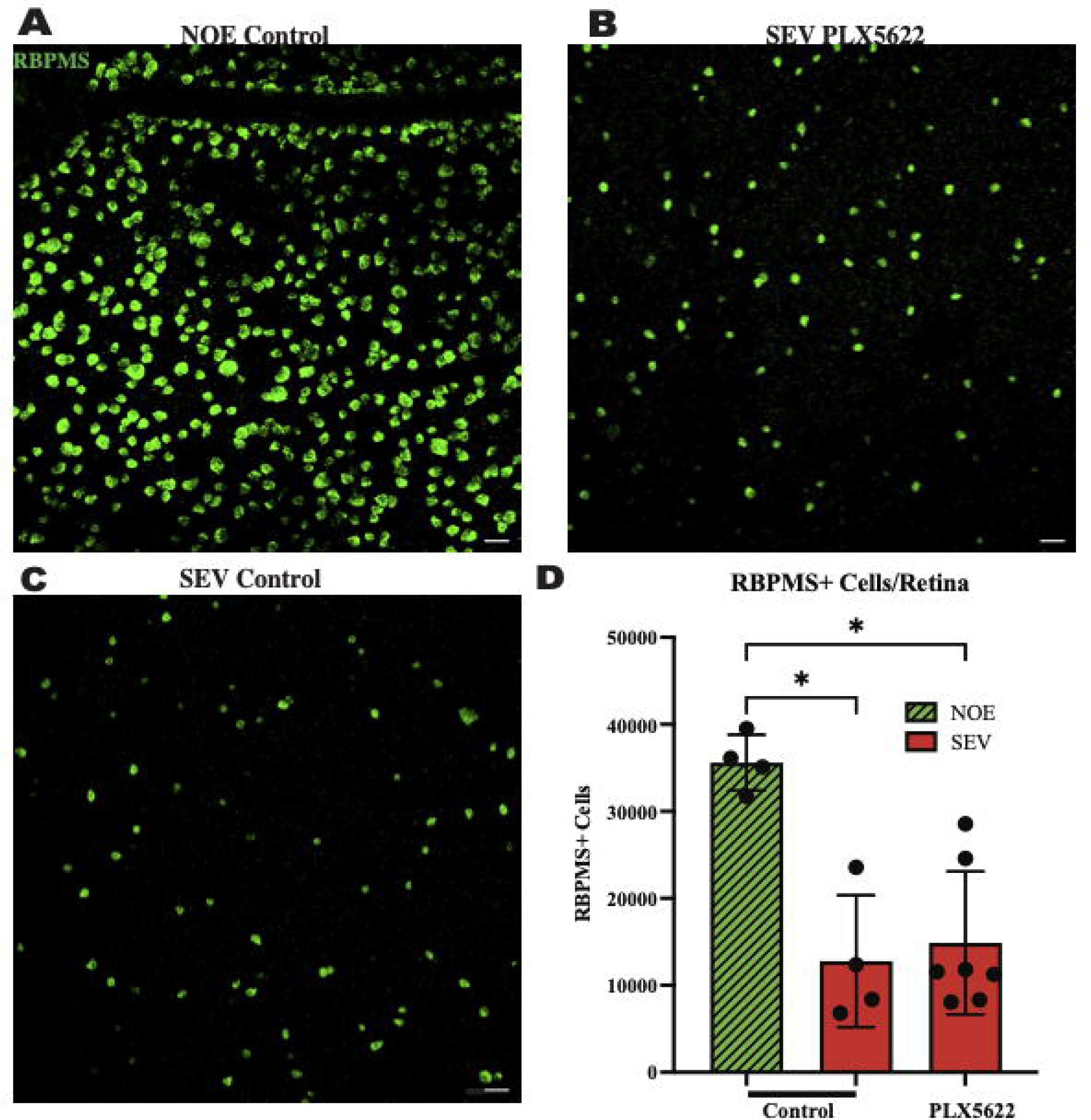
Reduction in microglia does not change RGC soma counts in severely damaged optic nerves. Representative images of RBPMS+ cells in retinal flatmounts of 12mos D2 retinas with No damage/early neurodegeneration (NOE) (**A**), severe damaged optic nerves on control diet (SEV.PLX5622l Diet) (**B**), and severe damaged optic nerves on PLX5622 diet (SEV.Control diet) (**C**). **D.** Graph of RBPMS+ cells per retina compared between damaged levels and treatment groups. NOE retinas (n=4) averaged 35597±3214 RBPMS+ cells. SEV control diet retinas (n=4) averaged 12778±7575 RBPMS+ cells, and SEV. PLX5622 diet retinas (n=6) averaged 14880±8206 RBPMS+ cells. Scale bar: 50µm. Data are presented as Mean±SEM. In **D**, Kruskal-Wallis test was performed.

## Discussion

It is unclear what role microglia have in regulating RGC death after injury, with reports suggesting they could have beneficial and detrimental (Howell et al., 2012, Heuss et al., 2018, Tribble et al., 2020, Howell et al., 2011, Rashid et al., 2019, Williams et al., 2019, Harder et al., 2020). To further understand the role of microglia in RGC death after an ocular hypertensive insult, we depleted microglia in D2 mice. We globally depleted microglia in D2 mice at 9.5 mos, before optic nerve damage occurs but after most D2 eyes develop elevated IOP (Anderson et al., 2005, Libby et al., 2005a). We found that global CSF1R inhibition via PLX5622 diet led to a robust reduction in microglia in retinal tissue. After 10 weeks on the PLX5622 diet, we did not see treatment-induced changes in IOP but did observe an increase in the prevalence of moderate-to-severe optic nerve damage. This suggests that microglia play a beneficial role in maintaining and protecting RGCs during an OHT injury. Similar results were observed in an acute model of OHT (Tan et al., 2022). We saw a much larger effect in terms of optic nerve damage in our aging OHT model with PLX5622 treatment compared to the acute inducible model (Tan et al., 2022). This may be because we introduced the PLX5622 diet after IOP elevation in many eyes, as opposed to depleting microglia prior to IOP elevation in the acute model of OHT study (Tan et al., 2022).

Global depletion of microglia appears damaging, suggesting an overall beneficial role of microglia in glaucomatous OHT. However, previous work by others showed that diminishing microglia activation via minocycline treatment (Bosco et al., 2008, Baptiste et al., 2005) promoted RGC survival in D2 mice, suggesting a damaging role for microglia. While minocycline has been suggested to promote neuroprotection in various injury models, it also affects other cells critical to inflammation such as peripheral macrophages (Dunston et al., 2011). Minocycline has been shown to enhance *in vitro* neuronal survival in the absence of microglia (Huang et al., 2010) further complicating the interpretation of the neuroprotective effect of minocycline in OHT. Moreover, previous work by us and others has suggested that targeting specific microglia-relevant molecular pathways, such as the complement cascade (Howell et al., 2011, Williams et al., 2016) or monocyte-like cell extravasation (Williams et al., 2019), provides neuroprotection. Taken together, the findings suggest nuanced roles for microglia in the context of glaucoma-relevant neurodegeneration. Understanding the differences between the damaging effects of global depletion of microglia, and the potential beneficial effects of targeting specific microglia-relevant processes will provide a more comprehensive view of the role of microglia in glaucoma.

Reduction of microglia through PLX5622 treatment has been shown to have beneficial effects in mouse models relevant to Alzheimer’s disease (Elmore et al., 2014, Spangenberg et al., 2019). In contrast, our work presented here and work by others (Tan et al., 2022) show PLX5622 is damaging in mouse models of glaucoma. These contrasting results suggest the mechanisms by which microglia respond to amyloid in Alzheimer’s disease (AD) to contribute to neuronal loss in the brain are different from microglia responses to ocular hypertension. In the face of injury or disease in both the brain and retina, microglia change from a homeostatic state to other transcriptional states, and whilst this transition from homeostatic microglia is thought to be initially beneficial, sustained inflammation can lead to neurotoxicity for RGCs (Rashid et al., 2019, Ramirez et al., 2017). In AD, the transition of homeostatic microglia to a disease-associated microglia phenotype is TREM2-dependent (Deczkowska et al., 2018). One study suggested that variations in TREM2 do not contribute to increased risk for primary open angle glaucoma (Margeta et al., 2020) while variations in TREM2 increased risk for AD. However, another study proposed TREM/TYROBP signaling may have a regulatory role in microglia responses in D2 mice (Tribble et al., 2020). Therefore, TREM2-dependent microglia responses may differ between glaucoma and AD, but this requires further investigation. Another microglia-relevant gene, apolipoprotein E (APOE), has been observed to have contradictory responses/affects between glaucoma and AD (Margeta et al., 2020). The E4 allele of *APOE* (*APOE4*) is the strongest genetic risk for Alzheimer’s disease, yet in glaucoma the presence of *APOE4* appears to be neuroprotective (Margeta et al., 2020). Determining the similarities and differences between microglia in diseases of the brain and the eye could help identify shared versus distinct therapeutic targets. In addition, examining the eye carefully when testing microglia-based treatments for AD will be important as treatments may have unexpected adverse effects in the eye.

Here, we focused on resident microglia, but infiltration of monocytes and monocyte-derived cells into the optic nerve head, the critical site of injury in glaucoma, has been previously been reported (Williams et al., 2019, Howell et al., 2012, Williams et al., 2017a). In the optic nerve head of D2 mice, monocytes were present prior to RGC axon damage but were missing in normotensive controls (Williams et al., 2019). Reduced extravasation of monocyte-like cells through genetic ablation of *Itgam*, (*Cd11b)* an important cell adhesion protein for tissue infiltration of monocytes, lead to lessened glaucomatous neurodegeneration in D2 mice, demonstrated through RGC axonal staining and soma counts (Williams et al., 2019). It is also possible that removing *Cd11b* decreases the damaging actions of microglia (Williams et al., 2019). Pharmacological inhibition of infiltrating monocytes into the retina/optic nerve prevented loss of RGC soma and axons (Williams et al., 2019). Further, inhibiting infiltration of monocytes cells into D2 optic nerve head through eye-specific radiation treatment protected the optic nerve from glaucomatous damage (Howell et al., 2012). On the other hand, significant glaucomatous damage occurred when monocyte-derived cells extravasated into the optic nerve head through deletion of *Glycam,* a negative regulator of extravasation, in neuroprotected irradiated mice (Williams et al., 2017a). Therefore, one possibility for the increased damage in PLX5622-depleted D2 mice is that the beneficial effects of resident microglia have been lost, enhancing the damaging effects of infiltrating monocytes.

In conclusion, our data showed that dietary PLX5622 significantly reduced microglia number and microglia gene expression in the retina of D2 mice. This ablation did not prevent RGC axon and soma loss. Rather, we saw an increase in the number of eyes with severe optic nerve damage, suggesting microglia play a critical neuroprotective role during glaucoma pathogenesis.

## Acknowledgments

We thank the JAX Flow Cytometry core facility for their expertise and assistance, Dr. Philipp Henrich at the JAX Microscopy core for training and assistance, and JAX Genomic Technologies for expert assistance. This study was supported by funding sources: EY035093 (GRH, RTL), Diana Davis Spencer Foundation (GRH), and by the University of Maine GSBSE T32 (CAD).

